# Androgen Deprivation-Induced TET2 Activation Fuels Prostate Cancer Progression via Epigenetic Priming and Slow-Cycling Cancer Cells

**DOI:** 10.1101/2025.03.26.645495

**Authors:** Lin Li, Siyuan Cheng, Yaru xu, Su Deng, Ping Mu, Xiuping Yu

**Affiliations:** Department of Biochemistry and Molecular Biology, LSU Health Shreveport, Shreveport, LA, 71130; Department of Urology, Yale University School of Medicine, New Haven, CT, 06511; Yale Cancer Center, Yale University School of Medicine, New Haven, CT, 06511

## Abstract

Advanced prostate cancer (PCa) frequently develops resistance to androgen deprivation therapy through various mechanisms including lineage plasticity. Slow-cycling cells (SCCs) have emerged as key players in adaptive responses to therapy, yet their role in PCa remains unclear. Through in silico analysis of single-cell RNA sequencing (scRNA-seq) data, we discovered that SCCs are enriched during pivotal stages of PCa progression, including the transition from androgen-dependent to castration-resistant states and the emergence of neuroendocrine PCa (NEPC). Using a tetracycline-inducible H2BeGFP reporter system, we confirmed SCC enrichment following androgen deprivation in both *in vitro* and *in vivo* models. Furthermore, we identified TET2 as a key regulator of SCCs, with its expression upregulated by androgen deprivation and positively correlated with SCC signature scores in PCa. Genome-wide 5-hydroxymethylcytosine (5hmC) profiling revealed increased hydroxymethylation after androgen deprivation, while TET2 knockdown reduced 5hmC levels at specific loci. Functional studies demonstrated that TET2 governs SCC maintenance, cell cycle progression, and DNA damage repair. Targeting TET2, either alone or in combination with an ATM inhibitor, significantly suppressed tumor growth, highlighting TET2 as a promising therapeutic target. Our study provides the first single-nucleotide resolution map of 5hmC dynamics in PCa, identifies a cell state driving epigenetic rewiring, and underscores the transformative potential of novel therapeutic strategies for advanced PCa.

## INTRODUCTION

Prostate cancer (PCa) is the most commonly diagnosed cancer in American men. While localized PCa can often be effectively treated with surgery or radiation therapy, managing advanced disease remains challenging. Androgen deprivation therapy (ADT) is the standard first-line treatment for advanced prostate adenocarcinoma (AdPCa)^1^. Although initial effective, ADT eventually fails, leading to the emergence of castration-resistant prostate cancer (CRPCa)^1^. Most CRPCa cases retain androgen receptor (AR) activity despite androgen deprivation through different mechanisms^1-6^. This finding has prompted the development of more potent second-generation AR antagonists, such as Enzalutamide (ENZ). While these therapies have shown efficacy in treating AR^+^ CRPCa^1,7^, some tumors evade AR-targeted treatments by adopting alternative phenotypes, including neuroendocrine prostate cancer (NEPCa, AR^-^/NE^+^), amphicrine PCa (AR^+^/NE^+^), and double-negative PCa (AR^-^/NE^-^)^8,9^. These phenotypic shifts are associated with poor prognosis and resistance to conventional therapies^7,8^.

Epigenetic reprogramming is one of the main contributors to the emergence of these alternative PCa phenotypes^10-12^. Among these mechanisms, DNA methylation plays a critical role, exhibiting distinct patterns between AdPCa and these alternative phenotypes^13-18^. The TET family of dioxygenases, including TET1, TET2, and TET3, catalyze the oxidation of 5-methylcytosine (5mC) to its derivatives, including 5-hydroxymethylcytosine (5hmC), 5-formylcytosine (5fC), and 5-carboxylcytosine (5caC), ultimately leading to DNA demethylation^19^. This dynamic regulation of the epigenome is essential for maintaining cellular plasticity and may contribute to the phenotypic shifts. Among the TET enzymes, TET2 has emerged as a key regulator of epigenetic reprogramming in cancer^20-22^.

A major contributor for cancer therapy resistance and relapse is the presence of slow-cycling cancer cells (SCCs)-a subset of stem-like, quiescent cells capable of re-entering the cell cycle and initiating tumor growth under favorable conditions^23-26^. SCCs have been identified in various malignancies, including breast, colorectal, renal, gastric cancer, melanoma, and glioblastoma^23-25^. Emerging evidence suggests that SCC abundance is strongly linked to treatment resistance^26^. Notably, elevated TET2 levels have been observed in therapy-resistant SCCs in colorectal cancer, glioblastoma and melanoma^26^. In these cancers, TET2 knockdown reduces SCC populations and increases apoptosis, highlighting its role in SCC survival and therapy resistance^26^. Recent studies suggest that TET2 is also involved in PCa therapy resistance and epigenetic reprogramming^20-22^. One study found that TET2 is expressed in treatment-resistant PCa cells, where it interacts with AR and its binding partner CXXC5 at noncanonical AR target loci, regulating gene expression^21^. Another study implicates TET2 in shaping the 5hmC landscape in PCa cells, contributing to lineage plasticity^22^. However, the specific role of TET2 in PCa SCCs and therapy resistance remains poorly defined.

In this study, we explored the role of TET2 in PCa progression and treatment resistance, focusing on its involvement in SCCs and DNA hydroxymethylation. We mapped 5hmC sites at single-nucleotide resolution in PCa cells and integrated these findings with comprehensive multi-omics analyses—including RNAseq and ChIPseq. Additionally, our research identified potential druggable targets and proposed combination therapy strategies to improve treatment efficacy for PCa. Our findings provide novel insights into the molecular mechanisms underlying SCCs, treatment resistance, and lineage plasticity in PCa.

## RESULTS

### 1. In silico analysis reveals SCC enrichment during key transition stages in PCa progression

Since SCCs are the potential population contributing to cancer treatment resistance and tumor relapse, a better understanding on the regulation and role of SCCs are important. Although the existence of SCCs has been shown in PCa, the detailed function and the regulation mechanism of SCCs in PCa is still understudied. To explore the role of SCCs in PCa progression, we first conducted an in silico analysis using single-cell RNAseq (scRNAseq) data from the mouse prostate single cell atlas (MoPSA)^9^. This dataset captures a continuous spectrum of PCa at various stages. By constructing a pseudotime trajectory, we mapped the progression of PCa from early-stage AdPCa to late-stage castrate-resistant AdPCa and ultimately to NEPCa (Figs. 1A & 1B).

**Figure 1.**
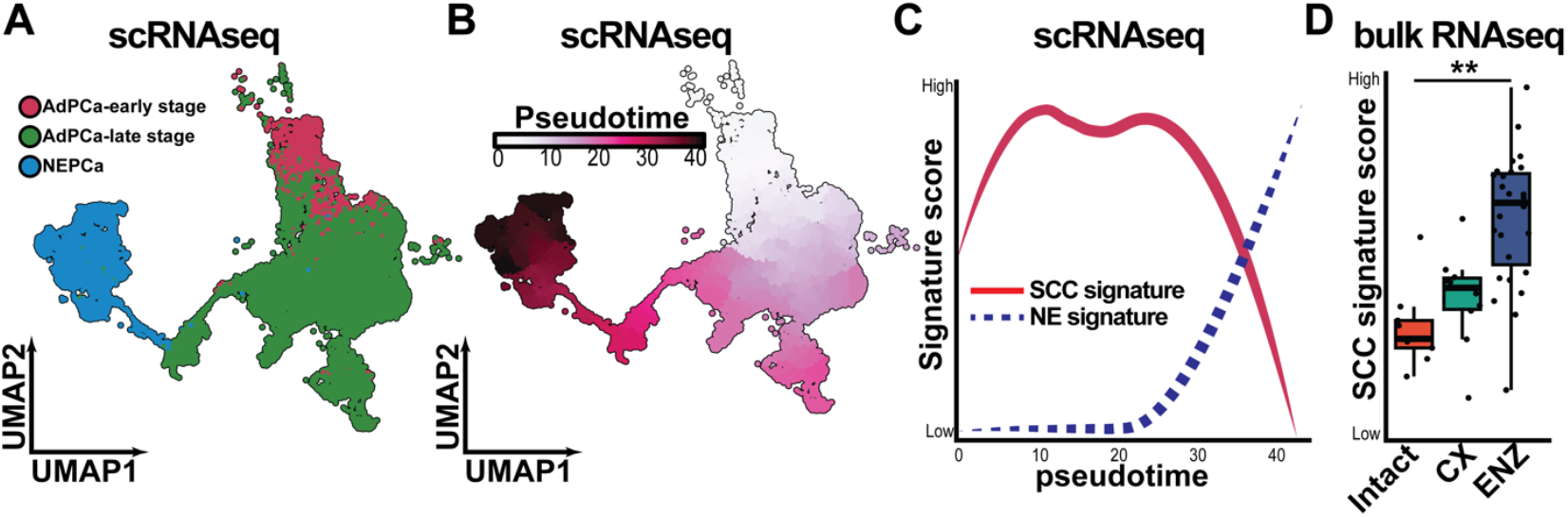
The SCC score is associated with PCa progression. (**A**) UMAP plot displaying distinct PCa cell populations within murine prostatic tumors based on scRNA-seq data from the MoPSA dataset. Pink: early-stage AdPCa; green: castrate-resistant AdPCa; blue: NEPCa. (**B**) Pseudotime trajectory predicting the predicted progression from early-stage AdPCa to castrate-resistant AdPCa and ultimately to NEPCa. The trajectory was reconstructed using UMAP embedding and the SimplePPT algorithm. Pseudotime values, calculated as the shortest path distances from a biologically defined root cell, represent relative progression rather than absolute time. (**C**) SCC score along the pseudotime trajectory of PCa progression. (**D**) SCC scores calculated from publicly available RNA-seq data (GSE211856) of LNCaP xenografts in intact mice, castrated (CX), and enzalutamide-treated (ENZ) mice. *Student’s t-test, p < 0*.*05*.

To assess SCC dynamics along this trajectory, we conducted single-sample Gene Set Enrichment Analysis (ssGSEA) using a previously defined SCC signature gene set^26^. Along the trajectory, we found that SCC scores progressively increased as PCa advanced from early to late-stage AdPCa, then to NEPCa, before ultimately decreasing in NEPCa cells (Fig. 1C). Notably, SCC scores exhibited two peaks: the first during the transition from early-to late-stage AdPCa (pseudotime ∼10), and the second during the transition from late-stage AdPCa to NEPCa (pseudotime ∼25), indicating SCC enrichment at these critical stages of PCa progression. To validate these findings, we examined SCC scores in bulk RNA-seq data (GSE211856) from LNCaP cell line-derived xenografts grown in intact, castrated, or enzalutamide (ENZ)-treated mice. Consistent with the scRNAseq result, SCC scores were significantly elevated in tumors from castrated and ENZ-treated mice, further supporting the association of SCCs with PCa progression and treatment resistance (Fig. 1D).

### 2. SCCs are enriched following androgen deprivation in PCa cells *in vitro* and *in vivo*

To further validate the presence and enrichment of SCCs in PCa, we utilized a tetracycline (Tet)-inducible histone 2B-eGFP (H2BeGFP) reporter system^26^ (Fig. 2A). This system labels cells with GFP upon doxycycline (DOX) treatment. Upon DOX withdrawal, the GFP intensity progressively dilutes with each cell division, allowing SCCs to be identified as cells that retain high GFP levels due to their slow proliferation.

**Figure 2.**
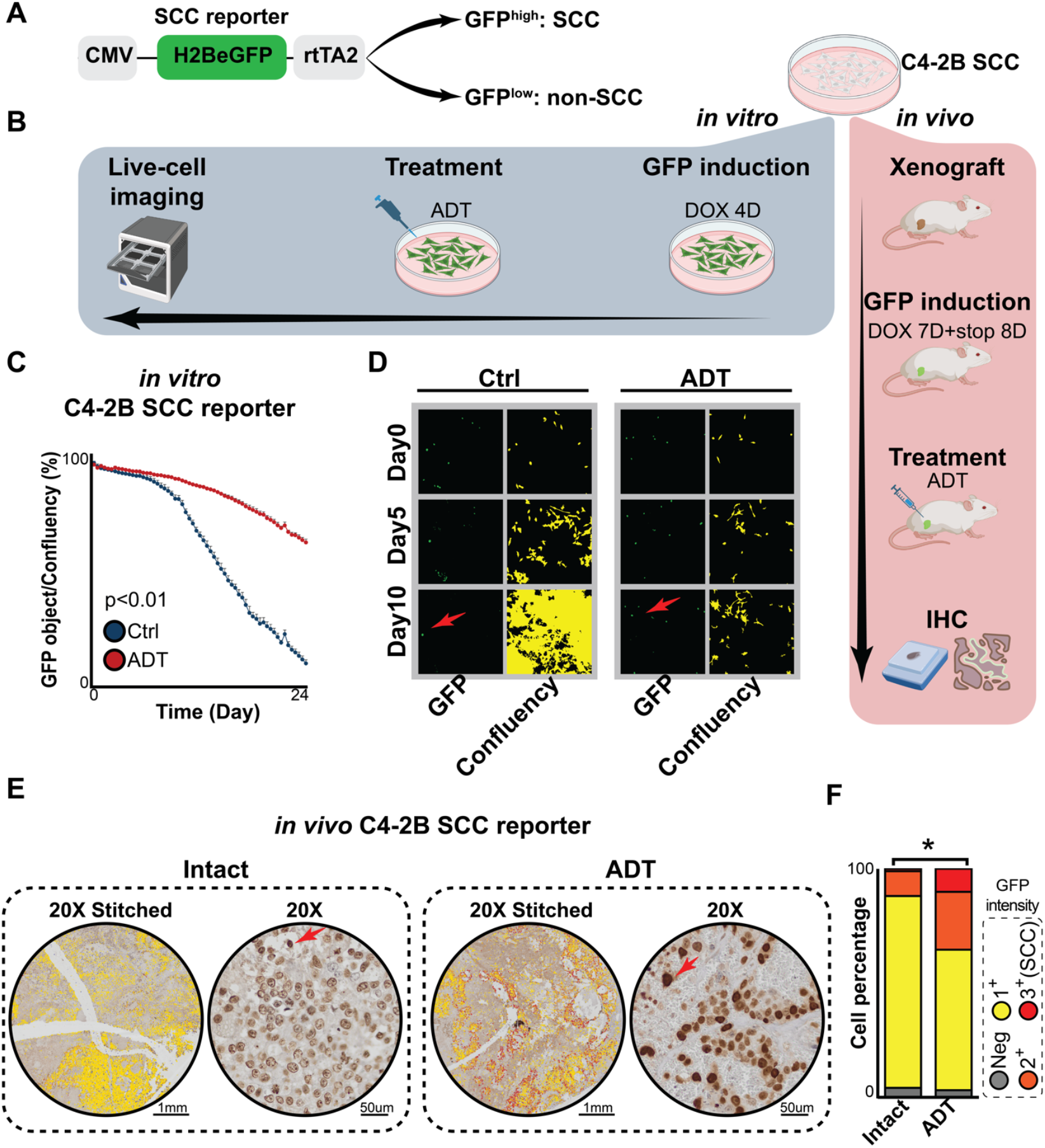
SCCs are enriched following androgen deprivation in PCa cells. (**A**) Schematic of the construct used for generating C4-2B SCC cells. The H2B-eGFP fusion gene expression is regulated by the reverse tetracycline transactivator (rtTA2) and activated by doxycycline (DOX). After DOX withdrawal, cells with high GFP intensity were considered as SCCs, while cells with low GFP intensity were non-SCCs Figure adapted from Addgene. (**B**) Experimental workflow for *in vitro* and *in vivo* SCC studies. *In vitro*, cells were treated with DOX for 4 days, followed by 10 days culturing with or without ADT. *In vivo*, xenografted tumors were labeled with DOX for 7 days. 8 days after DOX withdrawal, mice were either left untreated (Intact) or subjected to castration and ENZ (10 mg/kg) treatment (ADT) for 7 days. Workflow created using BioRender. (**C**) Quantification of GFP-positive cells under androgen-deprivation conditions compared to controls over a 10-day culture period, n=6. *Student’s t-test, p* < 0.01. (**D**) Live-cell imaging of GFP expression and confluency masks in control (Ctrl) and ADT groups at day 0, day 5, and day 10. Arrows indicate SCCs. (**E**) IHC staining of GFP in xenograft tumors. Upper panels show stitched images (20× magnification), and lower panels provide magnified views. Arrows highlight SCCs. Scale bars are indicated under the images. (**F**) Quantification of IHC results using QuPath. Cancer cells were categorized by GFP intensity into four groups: GFP-, GFP 1+ (0-0.2 raw pixel intensity, yellow color), GFP 2+ (0.2-0.4 raw pixel intensity, orange color), and GFP 3+ (0.4-0.8 raw pixel intensity, red color). Similar trends were observed in repeated experiments (n=3),two-way ANOVA, *p < 0*.*05*.

Using PCa C4-2B/SCC reporter cells, we conducted both *in vitro* and *in vivo* experiments (Fig. 2B). *In vitro*, reporter cells were treated with DOX for 4 days to induce GFP expression and then cultured under either control or androgen-deprived conditions (Fig. 2B). Live-cell imaging revealed that, in control group, GFP expression gradually diminished in most cells, but a subset retained high GFP levels, indicating the presence of pre-existing SCCs (Figs. 2C & 2D). Notably, under androgen deprivation, the proportion of GFP-high cells was significantly higher compared to control condition (Figs. 2C & 2D), indicating SCC enrichment in response to androgen deprivation.

*In vivo*, C4-2B/SCC reporter xenografts were established in mice. Mice were treated with DOX for 7 days to label PCa cells. Eight days after DOX withdrawal, mice were treated with or without castration (CX) and ENZ for seven days (Fig. 2B). The SCC population was assessed using immunohistochemistry (IHC) for GFP, and cancer cells were categorized into four groups based on GFP intensity: GFP^-^, GFP 1^+^, GFP 2^+^, and GFP 3^+^ groups (Fig. 2E). Since SCCs are defined as the less proliferative subset within the tumor, cells with the highest GFP intensity (GFP 3^+^) were classified as SCCs. Our analysis revealed that SCCs were present in tumors from both intact and castrated mice but were significantly enriched following androgen deprivation (Figs. 2E & 2F).

### 3. TET2 expression is enriched in SCCs and induced by androgen deprivation

Since SCCs were enriched following androgen deprivation, understanding their regulation could provide insights into PCa treatment resistance. Previous studies have identified TET2 as a key regulator of SCCs in colorectal cancer^26^. To investigate its association with SCCs in PCa, we first conducted a bioinformatics analysis using RNAseq data from ProAtlas^9^, a dataset integrating RNAseq profiles from hundreds human prostate specimens. Our analysis revealed a positive correlation between TET2 mRNA expression and SCC signature scores in PCa patient samples (Fig. 3A). This correlation was also observed in pan-cancer patient samples from The Cancer Genome Atlas (TCGA, Fig. 3B) and pan-cancer cell lines from the PCTA collection^27^ (Fig. 3C). To validate the in silico findings, we performed RT-qPCR to assess TET2 mRNA levels in SCC and non-SCC populations isolated from C4-2B/SCC reporter cells using fluorescence-activated cell sorting from both *in vitro* and *in vivo* experiments (Fig. 3D). TET2 expression was significantly higher in SCCs compared to non-SCCs in both settings (Figs. 3E & 3F).

**Figure 3.**
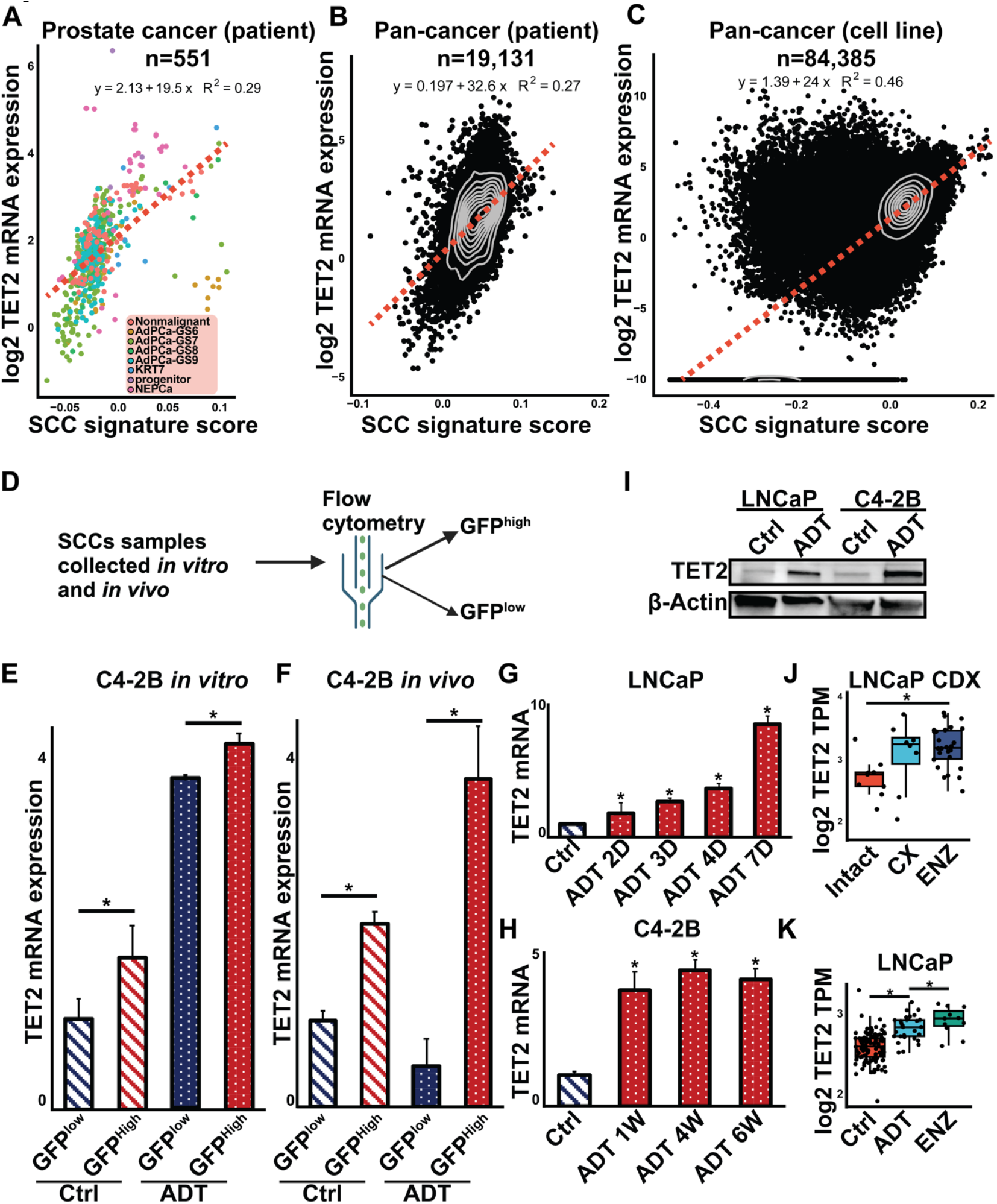
TET2 expression is higher in PCa SCCs and following androgen deprivation. (**A**) Correlation of TET2 expression with SCC score in PCa patient cohorts, including non-malignant tissues, Gleason scores 6 - 9, double-negative PCa (DNPCa), and NEPCa. (**B**) Correlation of TET2 expression with SCC score across TCGA pan-cancer patient cohorts. (**C**) Correlation of TET2 expression with SCC score in pan-cancer cell lines. (**D**) Workflow for isolating SCCs from non-SCCs using FACS to sort GFP^high^ (SCCs) and GFP^low^ (non-SCCs) cells. (**E**) RT-qPCR analysis of TET2 expression in GFP^high^ and GFP^low^ cells *in vitro* cultured with or without ADT. *Student’s t-test, p < 0*.*05*. (**F**) RT-qPCR analysis of TET2 expression in GFP^high^ and GFP^low^ cells *in vivo* xenograft from intact or castration + ENZ (ADT) mice. *Student’s t-test, p < 0*.*05*. (**G**) RT-qPCR analysis of TET2 expression in LNCaP cells cultured with or without ADT for 2, 3, 4, and 7 days, n=3. (**H**) RT-qPCR analysis of TET2 expression in C4-2B cells cultured with or without ADT for 1, 4 and 6 weeks, n=3. (**I**) Western blot analysis of TET2 protein levels in LNCaP and C4-2B cells with or without androgen deprivation. (**J**) The mRNA expression of TET2 in LNCaP xenografts in intact mice, castrated (CX), and enzalutamide-treated (ENZ) mice, data generated using publicly available RNA-seq data (GSE211856). *Student’s t-test, p < 0*.*05*. (**K**) Bioinformatic analysis of mRNA expression of TET2 in LNCaP cell cultured in full serum media, CS or ENZ conditions. *Student’ t-test, p < 0*.*05*.

Next, we investigated whether TET2 expression is regulated by AR signaling. TET2 mRNA levels were assessed in LNCaP and C4-2B cells cultured under control or androgen deprived conditions. Androgen deprivation significantly increased TET2 mRNA levels in both cell lines (Figs. 3G & 3H). This induction was also observed at the protein level, as TET2 protein expression was elevated following androgen deprivation (Fig. 3I). Consistent with these results, bioinformatics analysis of public datasets demonstrated that TET2 mRNA expression was significantly upregulated by androgen deprivation in both LNCaP-derived xenografts (GSE211856, Fig. 3J) and *in vitro*-cultured LNCaP cells (Fig. 3K). Further bioinformatics analysis of chromatin immunoprecipitation sequencing (ChIP-seq) data from LNCaP cells (collected in our recent study^28^) revealed AR binding sites in the *cis*-regulatory region of the TET2 gene, suggesting that AR directly regulates TET2 expression (sFig. 1). This finding provides a mechanistic explanation for the increased TET2 expression following androgen deprivation.

### 4. Genome-wide 5hmC levels increase in PCa cells following androgen deprivation

Given the established role of TET2 as a dioxygenase that oxidizes 5mC to 5hmC, we examined whether androgen deprivation-induced TET2 expression affects DNA hydroxymethylation levels in PCa cells. First, we performed enzyme-linked immunosorbent assay (ELISA) to assess global 5hmC changes following androgen deprivation. Our results revealed a significant increase in global 5hmC levels in LNCaP cells cultured under androgen deprivation conditions (Fig. 4A).

**Figure 4.**
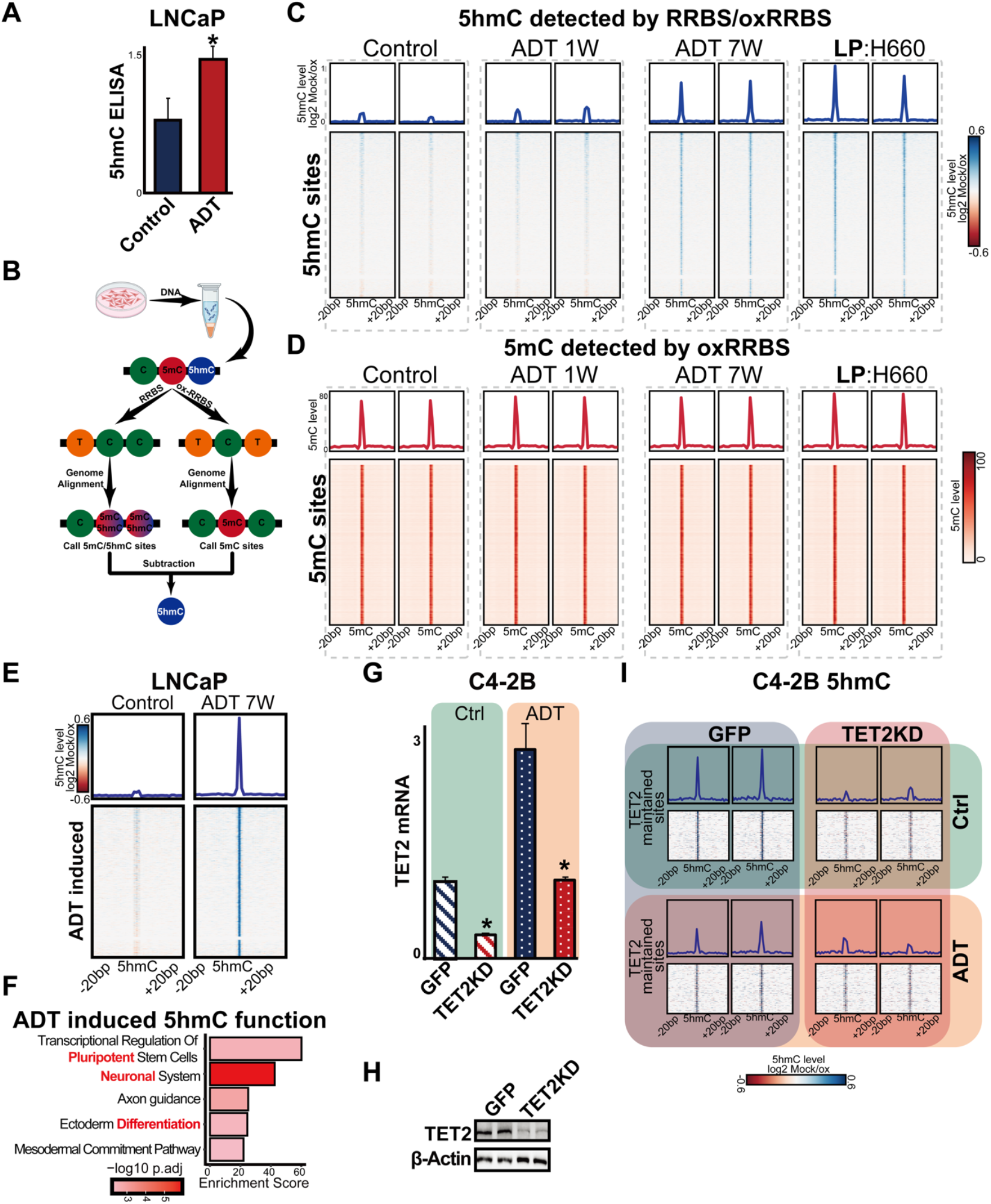
oxRRBS analysis on PCa cells treated with or without androgen deprivation. (**A**) 5hmC levels in LNCaP cells cultured in full serum media (Control) or CS (ADT) for 3 weeks assessed via ELISA, n=3. *Student’s t-test, p < 0*.*05*. (**B**) Workflow for RRBS and oxRRBS analysis. Control LNCaP cells and those cultured in full serum media (Control) or CS media (ADT) for 1 week, 7 weeks, and H660 cells were analyzed using oxRRBS assays, n=2. (**C**) Heatmap showing enriched regions for 5hmC across experimental conditions. (**D**) Heatmap showing enriched regions for 5mC in the same conditions as (**C**). (**E**) Heatmap showing differentially hydroxymethylated regions in LNCaP cells cultured with or without androgen depletion. (**F**) Enriched signaling pathways in differentially hydroxymethylated regions, highlighting associations with pluripotency, neuronal function, and ectoderm differentiation. TET2 knockdown efficiency was assessed by RT-qPCR (**G**) and western blot (**H**) in C4-2B cells. *Student t-test, p < 0*.*05*. (**I**) Heatmap showing 5hmC sites in C4-2B GFP or TET2KD cells cultured with or without ADT.

To identify specific 5hmC-enriched genomic sites altered by androgen removal, we conducted oxidative reduced representation bisulfite sequencing (ox-RRBS). This technique distinguishes between 5mC and 5hmC at single-nucleotide resolution by comparing RRBS (which detects both 5mC and 5hmC) with ox-RRBS (which selectively detects 5mC by converting 5hmC to 5fC) (Fig. 4B). Firstly, oxRRBS was performed in LNCaP cells treated with or without androgen deprivation as well as lineage plastic neuroendocrine prostate cancer (NEPCa) H660 cells. Remarkably, we observed a global increase in 5hmC levels in LNCaP cells following seven weeks of androgen deprivation (Fig. 4C). Notably, the 5hmC pattern in these cells closely resembled that of the NEPCa cells (Fig. 4C). However, this increase was only observed after seven weeks of androgen deprivation, not at one week, suggesting a gradual shift in epigenetic regulation over this period. In contrast, no significant global decrease in 5mC levels was observed (Fig. 4D), likely due to the relatively low abundance of 5hmC sites, which account for ∼1% of 5mC sites in the PCa genome observed from our oxRRBS data.

This global shift in 5hmC levels prompted us to investigate specific genomic regions undergoing hydroxymethylation changes. Genome-wide analysis identified 6,108 differentially hydroxymethylated sites in LNCaP cells cultured under androgen depletion condition for seven weeks (Fig. 4E). Functional enrichment analysis of genes near these sites revealed significant associations with pathways related to pluripotency, neuronal function, and ectoderm differentiation (Fig. 4F). Together, these findings suggest that androgen deprivation-induced 5hmC accumulation may contribute to lineage plasticity in PCa cells.

### 5. TET2 maintains a subset of 5hmC sites in PCa cells

To investigate the functional role of TET2 in PCa, we generated stable TET2 knockdown (TET2-KD) C4-2B and PC3 cells. The efficiency of TET2 knockdown was validated using RT-qPCR, Western blot and RNAseq analyses, confirming a significant reduction in TET2 expression (Figs. 4G, 4H & sFig. 2). As expected, androgen deprivation induced TET2 expression, while TET2 knockdown restored the expression to levels comparable to untreated C4-2B cells (Figs. 4G & 4H). Next, we conducted oxRRBS in C4-2B TET2-KD cells to assess the impact of TET2 knockdown on 5hmC levels. Our analysis revealed a reduction in 5hmC at a subset of hydroxymethylated sites under both control and androgen deprived conditions (Fig. 4I). These findings suggest that TET2 is essential for maintaining a subset of 5hmC sites in PCa cells, even when its basal expression is low in the presence of androgens.

### 6. TET2 regulates the expression of genes associated with cell-cycle progression and DNA damage response

To further investigate the molecular mechanisms of TET2 in PCa, we conducted RNA sequencing (RNA-seq) analysis on TET2-KD C4-2B and PC3 cells. The TET2 knockdown efficiency was further confirmed in PC3 and C42B cells (Fig. 5A). Differential expression analysis in C4-2B TET2-KD cells compared to C4-2B GFP control cells under androgen deprivation conditions identified 2,422 differentially expressed (DE) genes with an adjusted p-value < 0.05 (Fig. 5B). To explore the functional significance of TET2-regulated genes, we first examined SCC signature genes using ssGSEA. Heatmap showed that TET2 knockdown markedly reduced the SCC signature score in both PC3 and C4-2B cells. Notably, in C4-2B cells, while androgen deprivation elevated the score, TET2 knockdown reversed this effect, further supporting TET2’s critical role in maintaining SCCs in PCa (Fig. 5C).

**Figure 5.**
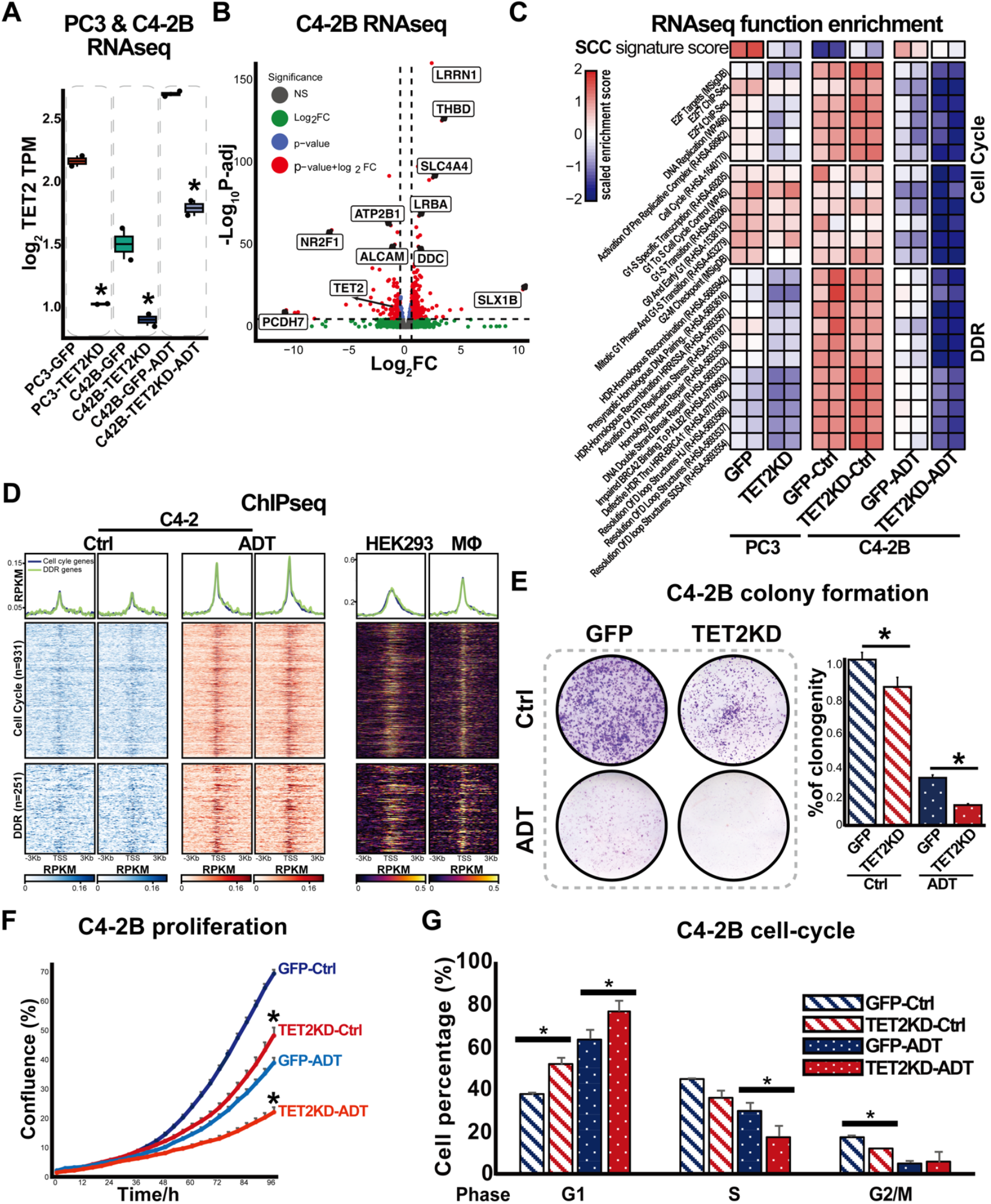
TET2 regulates the cell cycle in PCa cells. (**A**) TET2 mRNA levels in PC3 and C4-2B cells following TET2 knockdown, as determined by RNA-seq analysis. (**B**) Volcano plot showing differentially expressed (DE) genes in C4-2B control and TET2KD cells following androgen depletion. The fold-change and *p*-value were calculated between the two groups. All genes were shown, with only the most significant ones annotated with gene symbols. (**C**) Single-sample gene set enrichment analysis (ssGSEA) identifying signaling pathways affected by TET2 knockdown in PC3 and C4-2B cells. (**D**) Heatmap showing TET2 CHIPseq peaks in DNA damage response (DDR)- and cell cycle-related genes in C4-2 cells, under control and androgen deprivations. HEK293 and macrophage (Mφ) were used as positive controls. (**E**) Left: colony formation assay in C4-2B GFP or TET2KD cells cultured with or without ADT, n=6. Right: quantification of colony formation. *Student t-test, p < 0*.*05*. (**F**) Cell proliferation assay in C4-2B GFP or TET2KD cells cultured in full serum media (Ctrl) or CS + 20µM ENZ (ADT), n=6. measured using IncuCyte S3. (**G**) Cell cycle analysis of C4-2B GFP or TET2KD cells cultured with or without ADT, n=2. *Student t-test*, * *p < 0*.*05*.

Beyond SCC regulation, TET2 knockdown significantly altered gene signatures associated with two major functional categories: cell cycle progression and DNA damage response (DDR)-pathways (Fig. 5C). Re-analysis of publicly available TET2 ChIP-seq data further revealed that TET2 directly binds to numerous DE genes involved in cell cycle and DDR regulation (Fig. 5D), suggesting a direct transcriptional regulatory role. To validate the RNA-seq findings, RT-qPCR analysis was conducted to confirm the expression changes of selected DE genes related to cell cycle and DDR (sFig. 3).

### 7. TET2 regulates proliferation and cell cycle progression

Given that our RNAseq data revealed a role for TET2 in regulating cell cycle-related genes, we performed functional assays to further validate its impact on PCa cell proliferation and cell cycle progression. Firstly, cell proliferation and colony formation assays revealed a significant reduction in cell proliferation and colony formation in TET2-KD cells, under both androgen-sufficient and androgen-deprived conditions (Figs. 5E & 5F). Secondly, cell cycle analysis demonstrated that androgen deprivation induced G1-phase arrest, accompanied by a reduction in the S and G2/M phases. TET2 knockdown alone arrested cells in the G1 phase, with an even greater effect under androgen deprivations (Fig. 5G). Additionally, our RNAseq and qPCR analysis revealed that while most genes exhibited similar downregulation upon TET2 knockdown in both C4-2B and PC3 cells, certain key cell cycle-regulators, such as CDK1, PLK1, and PCNA, were downregulated only in C4-2B cells (sFig. 3) suggesting that TET2’s role in cell cycle regulation may be AR-dependent.

### 8. TET2 promotes DNA damage repair

Similarly, continued from our RNAseq data, to further confirm TET2’s role in DNA damage repair (DDR), we examined the protein levels of ATM, a key regulator of the DDR pathway, in C4-2B cells using Western blot analysis. Our results showed that both total ATM and phosphorylated ATM (p-ATM S1981) levels increased following androgen deprivation but decreased upon TET2 knockdown. (Fig. 6A). These findings suggest that DDR signaling is activated in response to androgen deprivation but attenuated when TET2 is depleted. To further confirm the TET2 maintained DDR in PCa cells, we directly counted the number of mutations from our C4-2B cells, comparing between control and TET2 knockdown conditions. The analysis revealed an increased number of mutations in cells under androgen deprivation, which was further exacerbated by TET2 knockdown (Fig. 6B). This suggests that TET2 loss contributes to the accumulation of DNA damage, particularly under androgen-deprived conditions. Eventually, we assessed phosphorylated H2AX (p-H2Ax) expression, a well-established marker of DNA double-strand breaks. Immunofluorescence staining revealed increased p-H2Ax accumulation in androgen-deprived C4-2B cells, with TET2 knockdown further exacerbating DNA damage (Fig. 6C left). Quantitative analysis confirmed significantly higher levels of DNA damage in TET2-knockdown cells, irrespective of androgen availability (Fig. 6C right).

**Figure 6.**
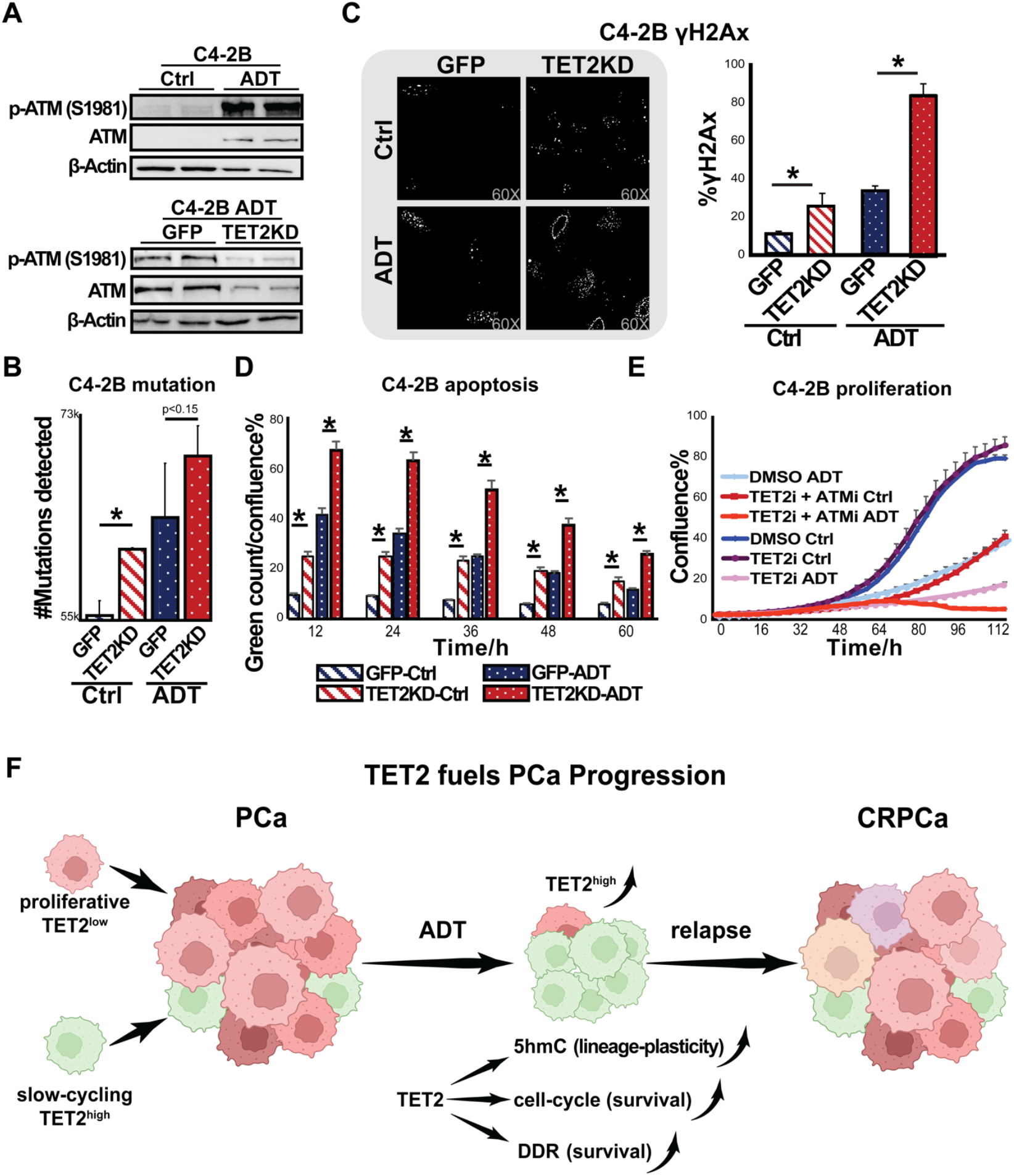
Knockdown of TET2 induces DDR and reduces cell proliferation. (**A**) Upper panel: Expression of ATM and p-ATM (S1981) in C4-2B cells cultured with or without ADT. Lower panel: Expression of ATM and p-ATM (S1981) in C4-2B GFP or TET2KD cells cultured with or without ADT. (**B**) Mutations analysis of RNAseq data from C4-2B GFP or TET2KD cells cultured with or without ADT, *Student t-test, * p <0* .*05*. (**C**) Left: Immunofluorescence staining of phospho-H2Ax (γH2AX) in C4-2B of C4-2B GFP or TET2KD cells cultured with or without ADT. Right: Quantification of γH2AX foci per cell (> 25 cells per group). *Student t-test, p < 0*.*05*. (**D**) Apoptosis analysis in C4-2B cells with or without TET2KD (Ctrl and TET2KD, respectively) cultured with or without ADT, n=6. Annexin V staining was used to detect apoptotic cells, followed by live-cell imaging for up to 60 hours. Apoptotic cells were quantified based on Annexin V-positive signals. *Student’s t-test, p < 0*.*05*. (**E**) Cell proliferation assay assessing the effects of TET2 and ATM inhibitors on PCa cell proliferation. C4-2B cells were cultured with or without ADT. treated with DMSO, Bobcat339 (TET2i, 10µM), AZD0156 (ATMi, 0.5µM), or a combination of both inhibitors, n=6. Live-cell imaging was conducted using Incucyte S3. (**F**) Proposed model depicting the role of TET2 in regulating PCa treatment resistance. The figure was generated using Biorender.

### 9. TET2 modulates apoptosis and represents a therapeutic target in PCa

Given the link between DNA damage accumulation and cell survival, we next examined whether TET2 influences PCa viability. Annexin V staining demonstrated that TET2 knockdown significantly increased apoptosis levels in both androgen-sufficient and androgen-deprived conditions, indicating that TET2 plays an essential role in supporting PCa cell survival (Fig. 6D).

As we observed the regulation of TET2 on ATM expression, DDR, and apoptosis, we hypothesized that co-targeting TET2 and ATM may have potential therapeutic implications. To test this hypothesis, we evaluated the effects of TET inhibitor Bobcat339 and observed a marked reduction in cell proliferation in androgen-deprived C4-2B cells. Notably, the combination of Bobcat339 with the ATM inhibitor AZD0156 exerted a synergistic effect, further enhancing growth suppression (Fig. 6E). These findings suggest that dual targeting of TET2 and ATM may offer a promising therapeutic approach for PCa treatment.

## DISCUSSION

Slow-cycling cells (SCCs) have been identified in many cancer types and are increasingly recognized for their crucial role in therapy resistance^26,29^. These cells exhibit distinct molecular and epigenetic characteristics that enable them to survive under stress conditions, such as androgen deprivation therapy in PCa. Understanding SCCs could provide key insights into the mechanisms that drive treatment resistance in PCa. In this study, we investigated the role of SCCs and TET2 in PCa progression and resistance to androgen deprivation.

Through in silico, *in vitro*, and *in vivo* SCC-reporter analyses, we identified SCCs in PCa and observed their enrichment following androgen deprivation. Notably, SCCs exhibited high TET2 expression, which was further enhanced by androgen deprivation. Functionally, TET2 knockdown reduced SCC-associated features, disrupted cell-cycle regulation, and impaired DNA damage response (DDR), leading to increased apoptosis and reduced proliferation in PCa cells. These findings establish TET2 as a key regulator of SCC maintenance, promoting DDR, and supporting cell survival under androgen-deprived conditions.

Based on these findings, we proposed a model in which SCC facilitate PCa survival under androgen deprivation conditions (Fig. 6F). While the majority of PCa cells undergo apoptosis following androgen deprivation, SCCs with high TET2 expression sustain survival by upregulating ATM and other DDR-related genes, thereby enhancing DNA damage repair and therapy resistance. This highlights TET2’s oncogenic role in driving resistance and positions SCCs as potential tumor-initiating populations, making them promising therapeutic targets. Further research is warranted to comprehensively characterize SCCs and elucidate the precise mechanisms by which TET2 sustains their survival.

Cancer lineage plasticity is closely associated with therapy resistance, enabling tumor cells to evade treatment by adopting alternative phenotypes^11^. Recent studies suggest that TET2 plays a critical role in epigenetic reprogramming and lineage plasticity in PCa^21,22^. Consistent with these findings, our study demonstrates that TET2 mediates 5hmC modifications. Given the established link between hydroxymethylation and DNA demethylation, TET2-driven 5hmC alterations may contribute to epigenetic reprogramming and therapy adaptation. While previous studies have examined the 5hmC landscape in PCa using 5hmC-seal sequencing^17,18,22,30^, the oxRRBS method employed in this study allows for the direct detection of both 5mC and 5hmC levels at single-nucleotide resolution, providing a more precise understanding of hydroxymethylation dynamics in PCa.

Additionally, we compared 5hmC levels between androgen-deprived AdPCa cells and the NEPCa cell line, H660. Notably, the 5hmC landscape in PCa cells following androgen deprivation closely resembles that of NEPCa cells, even when key NE transcription factors such as ASCL1 are not expressed. Furthermore. our recent research indicates that chromatin accessibility remains largely unchanged in PCa cells after androgen deprivation^28^, including within 5hmC-enriched regions. This suggests that rather than directly altering chromatin accessibility, the conversion of 5mC to 5hmC may serve as a priming mechanism, preparing these regions for future transcription factor binding, such as ASCL1, thereby facilitating trans-differentiation into NEPCa-like states.

In summary, we present the first single-nucleotide resolution map of 5hmC dynamics in PCa, highlighting TET2 as a central mediator of treatment resistance in PCa by regulating SCC maintenance, DDR activation, and lineage plasticity. Targeting TET2-ATM crosstalk offers a promising strategy to counteract SCC-driven resistance in advanced PCa.

## METHODS

### 1. Cell culture

LNCaP, C4-2B, and PC3 cells were obtained from ATCC and cultured in RPMI 1640 supplemented with 10% FBS, and 1% penicillin-streptomycin. H660 cells were cultured in ReproLife™ Reproductive Medium (Lifeline Cell Technology, #LL-0068). HEK293T cells were cultured in DMEM supplemented with 10% FBS, and 1% penicillin-streptomycin. To mimic androgen-deprivation condition, LNCaP and C4-2B cells were cultured in RPMI 1640 supplemented with 5% charcoal-stripped (CS)-FBS, or 5% charcoal-stripped FBS plus 20µM Enzalutamide (ENZ, #HY-70002, MedChemExpress).

### 2. Lentivirus transfection and transduction

Stable cell lines were generated using a lentiviral system. The SCCs reporter plasmid, which carries Doxycycline-inducible H2BeGFP expression (pSIN-TRE-H2BeGFP-rtTA2, #165494, Addgene), or TET2 knockdown construct (p-LV[shRNA]-mCherry:T2A:Puro-U6>hTET2 [shRNA#1], #VB900137-4927mqb, VectorBuilder) was used for transfection. Lentiviral particles were produced by transfecting HEK293T cells with the respective plasmids, along with pMD2.G, and psPAX2, using X-tremeGENE™ HP DNA Transfection Reagent (#6366244001, MilliporeSigma). The harvested viral supernatant was filtered with 0.45-µm filters (#25-246, Genesee Scientific) and used to transduce target cells in the presence of 4 µg/mL polybrene (#TR-1003-G, MilliporeSigma). Stable cells were selected with 4 µg/mL puromycin.

### *3. In vitro* and *in vivo* SCC quantification

For *in vitro* study, C4-2B SCCs reporter (C42B-SCC) cells were treated with 10 µg/mL Doxycycline (DOX) for 4 days, followed by cultured in normal FBS or CS-FBS + ENZ (20µM). SCCs were monitored via Incucyte live-cell imaging. GFP mask was set at a threshold of > 5 green calibrated unit (GCU). Fluorescence-activated cell sorting was conducted to separate SCCs (GFP high) from non-SCCs (GFP low) cells for RT-qPCR and RNAseq experiments.

For *in vivo* study, C42B-SCC cells were resuspended in PBS, mixed 1:1 with Matrigel, and subcutaneously injected into SCID mice. Once tumors were visible, mice received 0.5 mg/ml of DOX in drinking water for one week. Next, DOX was removed for 8 days, followed by treatment with or without castration plus ENZ (10 mg/kg) for 7 days. Mice were sacrificed, and tumors were collected as FFPE samples or dissociated into single cells for Fluorescence-activated cell sorting (FACS) analysis. All experiments were carried out in accordance with protocols approved by the Institutional Animal Care and Use Committee of LSU Health-Shreveport.

### 4. Immunohistochemistry (IHC) and immunofluorescence (IF) staining

IHC staining was performed using Vectastain elite ABC peroxidase kit (#PK-6100, Vector Laboratories) as described previously. GFP was detected using an anti-GFP antibody (#2956, dilution 1:400, Cell Signaling Technology) and visualized using DAB Substrate Kit (#SK-4100, Vector Laboratories). Tissue sections were counterstained, mounted, and imaged using a Zeiss microscope (White Plains). GFP intensity was quantified using QuPath v0.5.1. Based on the DAB OD mean value, intensity score was manually graded as 0 (negative, blue), 1^+^ (low, yellow, OD ≥0.2), 2^+^ (moderate, orange, OD ≥0.4), and 3^+^ (high, red, OD ≥0.8). For IF staining, primary antibodies included GFP (#sc-9996, dilution 1:100, Santa Cruz Biotechnology) and p-H2AX (#9718, dilution 1:200, Cell Signaling Technology). Images were captured using a Nikon fluorescence microscope (Melville, NY) and quantified using ImageJ.

### 5. FACS for SCCs sorting

For *in vitro* samples, DOX-induced cells were cultured in normal FBS or CS-FBS + ENZ (20µM) condition for 7 days before FACS analysis. A 3-day DOX induction was used as a positive control to isolate GFP^high^ SCCs.

For *in vivo* samples, dissociated tumor cells were resuspended in PBS. Live PCa cells were enriched using red blood cell lysis solution and EpCAM MicroBeads (#130-094-183 and #130-109-827, Miltenyi Biotec), following the manufacturer’s protocol. Cells were resuspended in PBS and subjected to FACS analysis.

### 6. RNA extraction, RT-qPCR, and RNA sequencing

All samples were lysed using TRI reagent and RNA was extracted using Direct-zol RNA Miniprep or Microprep kit (#R2053 and #R2060, Zymo Research) following the manufacturer’s protocol. cDNA was synthesized using ABScript III RT Master Mix for qPCR with gDNA Remover (#RK20429, Abclonal). qPCR was conducted using SYBR green PCR Supermix (#4344463, Applied Biosystem), with GAPDH as a normalization control. Primer sequences were listed in Table1.

### 7. Western blot

Cells were lysed in Laemmli SDS sample buffer and protein lysates were subjected to SDS-PAGE and Western blotting. Primary antibodies include beta-actin (sc-47778, dilution 1:500, Santa Cruz Biotechnology), TET2 (#45010, dilution 1:1000, Cell signaling Technology, #R1086-K1a, dilution 1:2000, Abiocode), ATM (#2873, dilution 1:1000, Cell signaling Technology, #A19650, dilution 1:1000, Abclonal), p-ATM at S1981 (#ab81292, dilution 1:1000, Abcam). HRP-conjugated secondary antibodies (#AS014, and #AS003, dilution 1:5000, Abclonal) were used. Protein bands were visualized using Tanon™ High-sig ECL Western Blotting Substrate (#180-5001, Abclonal) and Chemidoc™ Touch Imaging System (Bio-Rad).

### 8. Apoptosis analysis using Annexin V staining

Cells were seeded in 96-well plate overnight, then cultured in normal FBS or CS-FBS + ENZ (20 µM). Apoptosis was assessed using 10 µL/mL Annexin V-Alexa Fluor 488 Conjugates (#A13201, Invitrogen), with FITC+ cells quantified via Incucyte S3 live-cell imaging.

### 9. Cell proliferation and colony formation

For proliferation assay, 1,000 cells were seeded per well in 96-well plates, treated accordingly, and monitored via Incucyte S3 live-cell imaging. Colony formation assays were conducted by seeding 1000 cells per well in 6-well plates. After 2 weeks of treatment, colonies were fixed with 70% ethanol and stained with 0.5% crystal violet. After scanning the plate, the colonies were quantified by dissolving the stain in 2-ethoxyethonal followed by absorbance measurement at OD560.

### 10. DNA extraction and ELISA

Genomic DNAs from LNCaP cells cultured with normal FBS or CS-FBS for different time points were extracted using E.Z.N.A.® Tissue DNA Kit (#D3396, Omega Bio-tek). Then 100ng DNA was used to perform 5-hmC ELISA using MethylFlash Global DNA Hydroxymethylation (5hmC) ELISA Easy Kit (Colorimetric) (#P-1032, Epigentek) following the manufacturer’s instructions. The absorbance was read using microplate reader at OD450 nm.

### 11. oxRRBS

Bisulfite sequencing and oxidative bisulfite sequencing were conducted using Ovation® RRBS/oxBS kit (#0553, Tecan) following the manufacturer’s instruction. Then the prepared libraries were sequenced by Novogene. The nf-core methylseq pipeline was used for sequencing alignment and methylation calls extraction. Specifically, the bowtie2 was used as aligner and GRCh38 was used as reference genome. The Bismark was used for downstream analysis. The 5hmC sites were identified by contrasting the mock and ox samples using calculateDiffMeth function from “methylKit” R package (https://github.com/al2na/methylKit) with “difference=50, qvalue=0.05” as cutting off condition. In addition, the deeptools (https://github.com/deeptools/deepTools) was used for visualization.

### 12. Cell cycle analysis

Cells were harvested, washed and suspended in PBS, and fixed in 70% ethanol. Fixed cells were treated with 50 µg/mL propidium iodide (PI, #AAJ66764MC, Fisher Scientific) and 0.5 µg/mL RNase A, followed by flow cytometry analysis.

### 13. RNAseq data library preparation and processing

Total RNA was sequenced uisng NovaSeq X Plus Series (150 bp pair-end reads). Data preprocessing, alignment, and gene expression analysis were performed using FastQC, Trim Galore, STAR, and Salmon. Differential expression analysis was conducted using DESeq2, and gene signature activity scores were calculated using singscore. Besides, the nf-core rnavar pipeline was used for RNAseq data based variant calling analysis.

## Acknowledgement

This work was supported by NIH R01 CA226285, Louisiana State University Health Sciences Center at Shreveport FWCC Stimulus award, and LSU Collaborative Cancer Research Initiative funding to X. Yu. LSU Health Shreveport FWCC Carroll Feist predoctoral Fellowship to L. Li. We would like to thank the INLET high-throughput imaging core at LSUHSC-Shreveport (RRID: SCR_024990) for Incucyte assistance.

## Figure legend

**Supplementary Figure 1.**
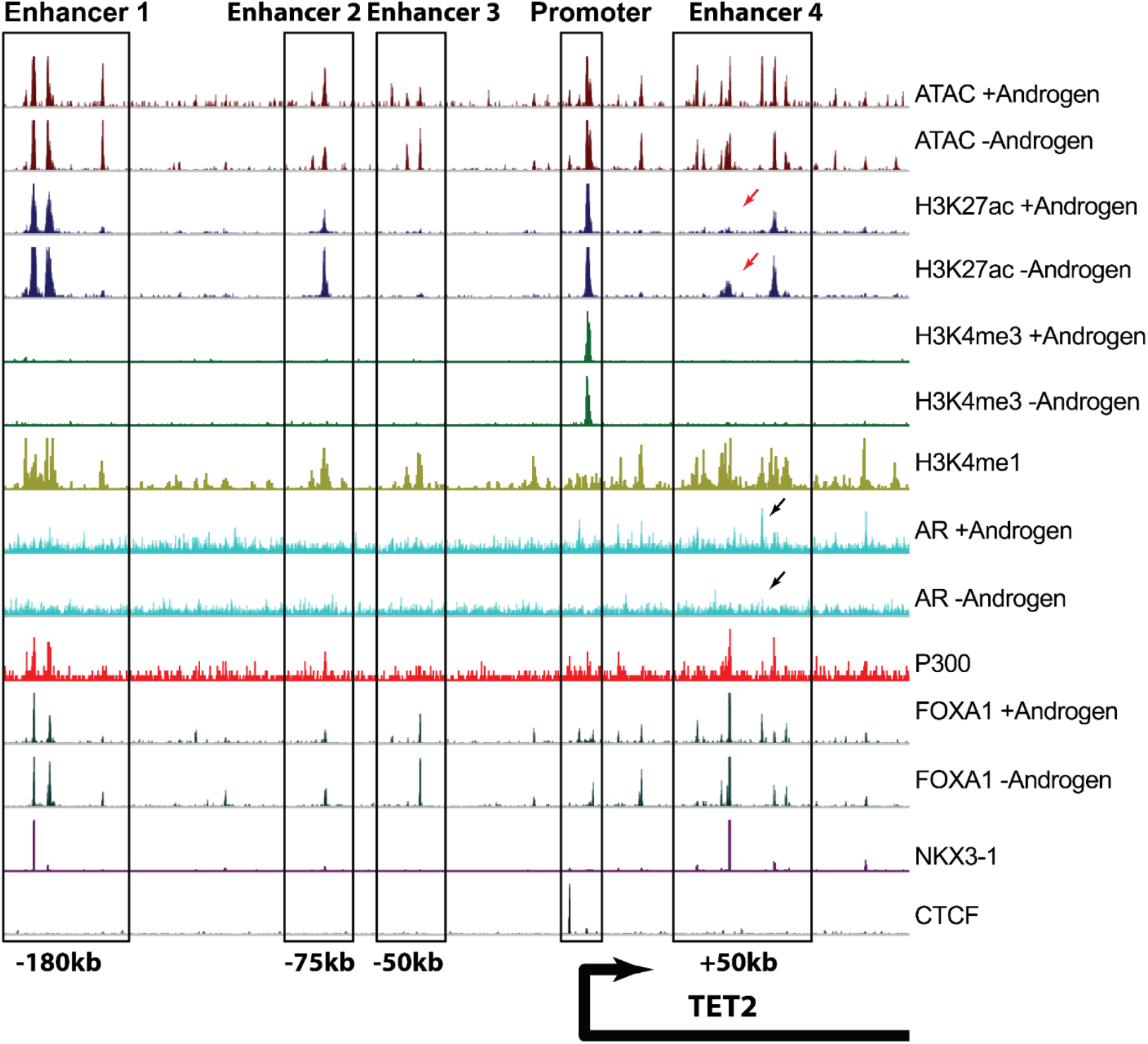
AR directly binds to the cis-regulatory sequence of TET2. CHIPseq and ATACseq data from LNCaP cells cultured with or without androgen illustrating distinct binding patterns of various factors at the TET2 locus. Red arrows indicate increased H3K27ac peaks following androgen deprivation, suggesting activation of this region. Blue arrows indicate reduced AR binding peaks after androgen removal.

**Supplementary Figure 2.**
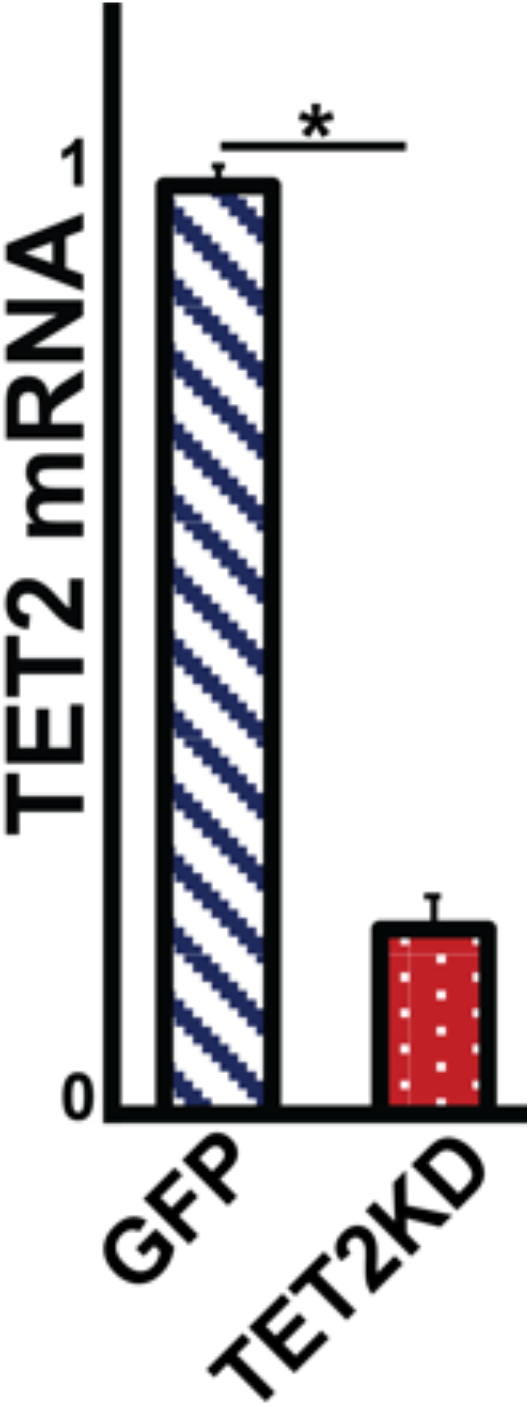
Validation of TET2 knockdown efficiency in PC3 cells. TET2 knockdown efficiency was confirmed by RT-qPCR in PC3 cells. *Student t-test, p < 0*.*05*.

**Supplementary Figure 3.**
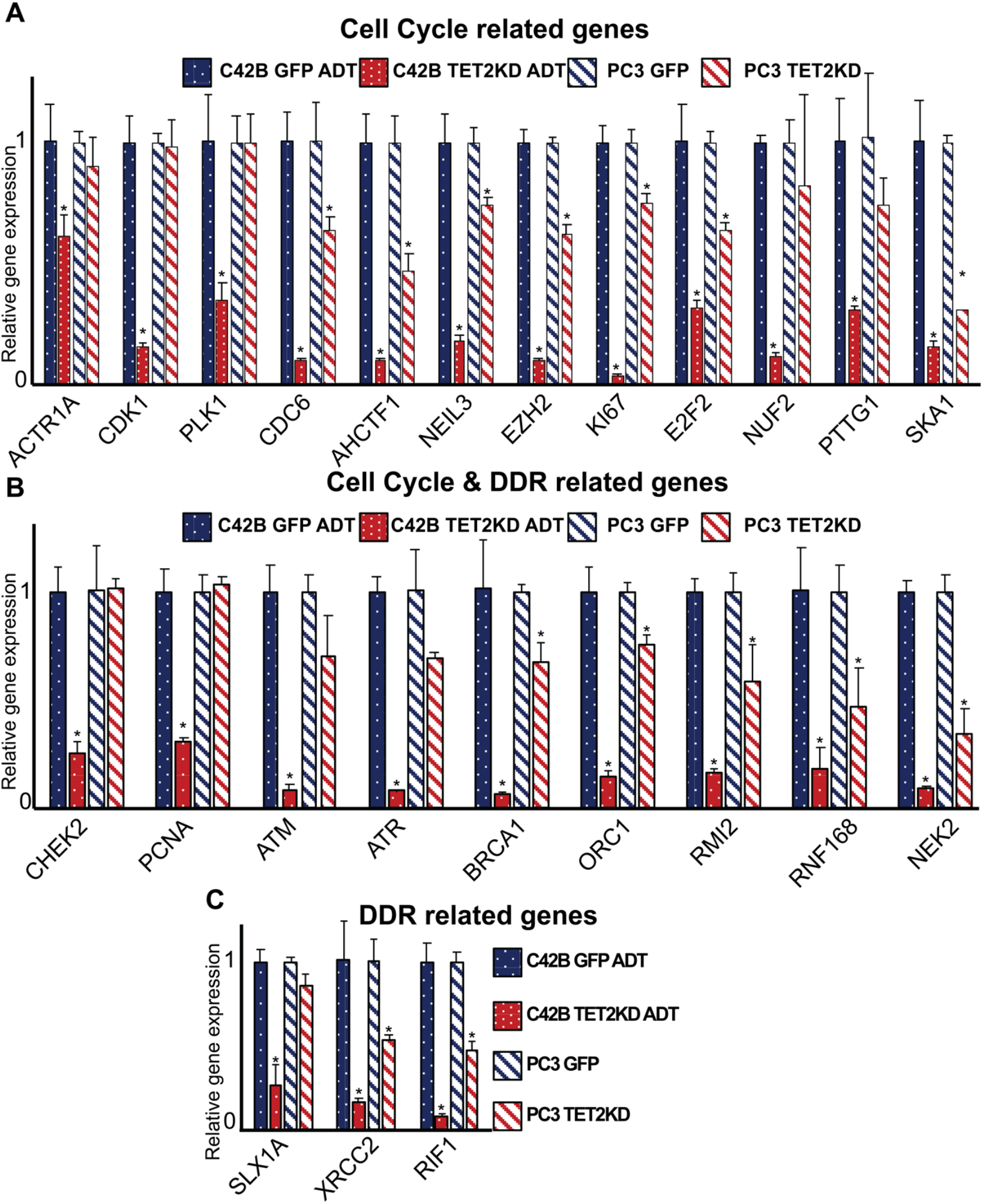
TET2 regulates the expression of cell cycle- and DDR-related genes. RT-qPCR analyses of cell cycle-related genes in PC3 GFP and TET2KD cells cultured in full serum media, C4-2B GFP and TET2KD cells cultured in ADT conditions, n=3. *Student t-test, p < 0*.*05*.

## REFERENCES

1 Debes, J. D. & Tindall, D. J. Mechanisms of androgen-refractory prostate cancer. New England Journal of Medicine 351, 1488–1490 (2004).

2 Li, X. & Mu, P. Restoring our ubiquitination machinery to overcome resistance in cancer therapy. Oncoscience 11, 43 (2024).

3 Li, X. & Mu, P. The critical interplay of CAF plasticity and resistance in prostate cancer. Cancer research 83, 2990–2992 (2023).

4 Blatt, E. B. et al. Overcoming oncogene addiction in breast and prostate cancers: a comparative mechanistic overview. Endocrine-related cancer 28, R31–R46 (2021).

5 Rodriguez Tirado, C. et al. UBE2J1 is the E2 ubiquitin-conjugating enzyme regulating androgen receptor degradation and antiandrogen resistance. Oncogene 43, 265–280 (2024).

6 Li, X. et al. Loss of SYNCRIP unleashes APOBEC-driven mutagenesis, tumor heterogeneity, and AR-targeted therapy resistance in prostate cancer. Cancer Cell 41, 1427–1449 (2023).

7 Tran, C. et al. Development of a second-generation antiandrogen for treatment of advanced prostate cancer. Science 324, 787–790 (2009).

8 Vlachostergios, P. J., Puca, L. & Beltran, H. Emerging variants of castration-resistant prostate cancer. Current oncology reports 19, 1–10 (2017).

9 Cheng, S. et al. Unveiling novel double-negative prostate cancer subtypes through single-cell RNA sequencing analysis. npj Precision Oncology 8, 171 (2024).

10 Davies, A. H., Beltran, H. & Zoubeidi, A. Cellular plasticity and the neuroendocrine phenotype in prostate cancer. Nature Reviews Urology 15, 271–286 (2018).

11 Beltran, H. et al. The Role of Lineage Plasticity in Prostate Cancer Therapy ResistanceLineage Plasticity in Prostate Cancer Resistance. Clinical cancer research 25, 6916–6924 (2019).

12 Wen, P. et al. Hyd/UBR5 defines a tumor suppressor pathway that links Polycomb repressive complex to regulated protein degradation in tissue growth control and tumorigenesis. Genes & Development 38, 675–691 (2024).

13 Donkena, K. V., Young, C. Y. F. & Tindall, D. J. Oxidative stress and DNA methylation in prostate cancer. Obstetrics and gynecology international 2010, 302051 (2010).

14 Kobayashi, Y. et al. DNA methylation profiling reveals novel biomarkers and important roles for DNA methyltransferases in prostate cancer. Genome research 21, 1017–1027 (2011).

15 Beltran, H. et al. Divergent clonal evolution of castration-resistant neuroendocrine prostate cancer. Nature medicine 22, 298–305 (2016).

16 Zhao, S. G. et al. The DNA methylation landscape of advanced prostate cancer. Nature genetics 52, 778–789 (2020).

17 Gallon, J. et al. DNA methylation landscapes of prostate cancer brain metastasis are shaped by early driver genetic alterations. Cancer research 83, 1203–1213 (2023).

18 Franceschini, G. M. et al. Noninvasive Detection of Neuroendocrine Prostate Cancer through Targeted Cell-free DNA Methylation. Cancer discovery 14, 424–445 (2024).

19 Ito, S. et al. Tet proteins can convert 5-methylcytosine to 5-formylcytosine and 5-carboxylcytosine. Science 333, 1300–1303 (2011).

20 Takayama, K.-i. et al. TET2 repression by androgen hormone regulates global hydroxymethylation status and prostate cancer progression. Nature communications 6, 1–16 (2015).

21 He, Y. et al. A noncanonical AR addiction drives enzalutamide resistance in prostate cancer. Nature communications 12, 1521 (2021).

22 Xu, Y. et al. ZNF397 Deficiency Triggers TET2-Driven Lineage Plasticity and AR-Targeted Therapy Resistance in Prostate Cancer. Cancer Discovery 14, 1496–1521 (2024).

23 Roesch, A. et al. A temporarily distinct subpopulation of slow-cycling melanoma cells is required for continuous tumor growth. Cell 141, 583–594 (2010).

24 Chen, J. et al. A restricted cell population propagates glioblastoma growth after chemotherapy. Nature 488, 522–526 (2012).

25 Kreso, A. et al. Variable clonal repopulation dynamics influence chemotherapy response in colorectal cancer. Science 339, 543–548 (2013).

26 Puig, I. et al. TET2 controls chemoresistant slow-cycling cancer cell survival and tumor recurrence. The Journal of clinical investigation 128, 3887–3905 (2018).

27 Cheng, S., Li, L. & Yu, X. PCTA, a pan-cancer cell line transcriptome atlas. Cancer Letters 588, 216808 (2024).

28 Li, L., Cho, K. H., Yu, X. & Cheng, S. Systematic multi-omics investigation of androgen receptor driven gene expression and epigenetics changes in prostate cancer. Computers in Biology and Medicine 189, 110000 (2025).

29 Lee, H. et al. Cell cycle arrest induces lipid droplet formation and confers ferroptosis resistance. Nature communications 15, 79 (2024).

30 Sjöström, M. et al. The 5-hydroxymethylcytosine landscape of prostate cancer. Cancer research 82, 3888–3902 (2022).

